# CryoDDM: CryoEM denoising diffusion model for heterogeneous conformational reconstruction

**DOI:** 10.64898/2025.12.10.693455

**Authors:** Fuwei Li, Yuanbo Chen, Hao Dong, Chenxuan Ji, Xinsheng Wang, Chuanyang Zhang, Zupeng Wang, Bin Hu, Fa Zhang, Xiaohua Wan

**Affiliations:** Key Laboratory of Brain Health Intelligent Evaluation and Intervention, Beijing Institute of Technology, Beijing, 100081, China; School of Medical Technology, Beijing Institute of Technology, Beijing, 100081, China; WQ & UCAS Research Academy Intelligent Computing Center (WRA-ICC), China

**Author notes:** Contributing authors.

**Keywords:** Cryo-EM, denoising diffusion model, 3D classification, heterogeneity conformation

## Abstract

Heterogeneous protein reconstruction can reveal the relationship between protein dynamics and function, which represents one of the foremost challenges in cryogenic electron microscopy (cryo-EM) single-particle analysis (SPA). However, high-intensity noise results in inaccurate parameter estimation during classification and reconstruction, making it difficult to capture the subtle motions of proteins. Here, we present the Cryo-EM denoising diffusion model (CryoDDM), which denoises images while preserving high-frequency structural information, benefiting conformational heterogeneity classification and reconstruction. The image denoised by CryoDDM can be used to improve downstream analysis, enhance reconstruction resolution, discover new protein conformations, and map out conformational variations. Validated against tree diverse experimental datasets, spanning a proteasome, a membrane protein and a spike protein, our results consistently demonstrate the accuracy and superiority of CryoDDM. By guiding reconstruction with denoised images, CryoDDM enables high-resolution reconstruction of heterogeneous conformations, paving the way to illuminate fundamental questions in structural biology. The CryoDDM project are published at https://github.com/BIT-FuweiLi/CryoDDM.

## 1 Introduction

The dynamic three-dimensional (3D) structures of proteins are closely related to the life processes they govern^1^. Method that unraveling protein conformational changes can provide a better understanding of protein activity and enhance drug design capabilities. In recent years, cryogenic electron microscopy (Cryo-EM) single-particle analysis (SPA) algorithms have been used to reconstruct many high-resolution 3D structures of biological proteins^2, 3^. However, the differences between different conformations are minimal, and high-intensity noise can introduce errors in the similarity calculations during the whole reconstruction pipeline. When the error exceeds the conformation differences, the algorithm will no longer effectively distinguish structural differences between the conformations. As a result, particle images in different conformations become mixed, often requiring additional post-processing to extract their heterogeneous characteristics. Therefore, high noise plays a crucial role in dynamic protein classification and reconstruction.

Manifold embedding^4^, multi-body refinement^5^, 3DVA^6^ and Zernike3D^7^ are represented an early attempt to reveal continuous molecular motions. They often assume that proteins follow certain structural patterns, random flexible motion, polytope binding, spherical twisting. Based on these assumptions, they construct mathematical models to better understand the structural heterogeneity of proteins. However, they often require accurate prior information and are prone to artifacts during the motion process. Deep learning can capture the conformation heterogeneity directly from the data itself. CryoDRGN^8^ and cryoDRGN2^9^ use variational autoencoder^10^ (VAE) to reveal the heterogeneous states using Fourier domain feature. CryoNeFEN^11^ employs a similar methodology to characterize protein heterogeneity directly in real space. 3DFlex ^12^ employs deformation fields to model protein motion, whereas Dynamight^13^ leverages the estimated motion fields to inversely align protein projections, thereby enhancing the resolution of reconstructed protein structures. CryoStar ^14^, on the other hand, incorporates an atomic model and introduces a set of collision constraints to the atomic framework, aiming to mitigate erroneous distortions and prevent atomic overlaps during the estimation of point cloud motions.

These methods are post-processing techniques for classification and reconstruction that do not account for heterogeneity throughout the entire reconstruction pipeline and rely on high-quality reconstruction results. However, high-intensity noise in cryo-EM images significantly affects the accuracy of parameter estimation during classification and reconstruction^15–18^, making it difficult to achieve high-quality reconstruction results^19, 20^ and stabilize the outcomes.

CryoSPARC^18^ and RELION^17, 21^ use filtering^22^ and masking^23, 24^ algorithms to reduce noise interference during reconstruction alignment. However, these algorithms cannot distinguish accurately between noise and structural information, sacrificing denoising performance to preserve structural details. Topaz^25^, based on the Noise2Noise^26^ framework, introduces a pre-trained denoising model that significantly enhances the SNR of cryo-EM images. Nevertheless, it tends to damage structural information, limiting its use primarily to particle picking. NT2C^27^ and Mscale^20^ attempt to preserve high-frequency information through amplitude correction after denoising, although they have limitations in ensuring the authenticity of the recovered structural information. Existing denoising methods cannot distinguish accurately between noise and structural information, significantly limiting their applicability and causing classification and reconstruction to remain affected by high-intensity noise. As a result, achieving high-quality classification and reconstruction becomes challenging, hindering accurate heterogeneous conformational analysis.

Here, we propose the Cryo-EM denoising diffusion model (CryoDDM) to highlight the heterogeneity of protein conformations. Compared to other post-processing methods that rely on high-precision 3D reconstruction results, our approach addresses the noise interference directly in the whole reconstruction pipeline, thereby obtaining more accurate and high-precision heterogeneous protein structures. We leverage the interpretability of the diffusion model to guide the neural network in distinguishing noise from structural information, achieving denoising while preserving structural information at near-atomic resolution and enhancing image quality. Denoised images overcome high-intensity noise interference, enabling more accurate parameter estimation throughout the single-particle-analyis(SPA) whole reconstruction pipeline, such as particle picking, 2D/3D classification and reconstuction, and leading to improved reconstruction result, which is particularly useful for heterogeneous conformational analysis.

We validate the CryoDDM and CryoDDM-based reconstruction pipeline on 6 public experimental datasets with different protein structural feature, achieving the most variable conformations. The EMPIAR-12087 dataset^28^, consisting of cryo-EM micrographs of human RYBP–PRC1 bound to an H2Aub1-modified mononucleosome, depicts a disc-like DNA–protein complex. Through CryoDDM, we reveal the dynamic behavior of the RYBP–Ub module within this assembly. We demonstrate this ability with a dataset of TRPV1 ion-channel particles (EMPIAR-10059^29^), and recover the motions that generally agree with other methods^12, 14^, further identifying specific movement sites. CryoDDM performs well on highly symmetric cylindrical proteins, T20S proteasome^30^,revealing the overall motion of the protein-binding sites and providing structural evidence that completes previous speculations regarding the open and closed states of the *α*-subunit N-terminus in the 20S proteasome channel^31^. Finally, we recover *α*V*β*8 integrin (EMPIAR-10345^32^),,significantly improving the reconstruction resolution and accuracy. Our experiments demonstrate that CryoDDM denoises images while preserving structural information, resulting in higher-quality images, more accurate parameter estimation, and ultimately improved reconstruction, which is particularly useful for heterogeneous conformational analysis.

## 2 Results

### 2.1 CryoDDM-based reconstruction pipeline for conformational analysis

Existing heterogeneous conformation reconstruction pipelines primarily include motion correction^33^, CTF correction^34, 35^, denoising, particle picking, 2D classification, 3D classification, homogeneous refinement, local refinement, and heterogeneity analysis methods such as 3DFlex^12^. However, current denoising method struggle to balance noise removal with the preservation of structural information. Thus, denoised images ^25, 36, 37^ can only be used for particle picking, indicating that high-intensity noise still significantly affects subsequent processes.

CryoDDM preserves the structural information of the particles during the denoising process, making the denoised images available for optimizing the estimation of parameters in subsequent heterogeneous conformation reconstruction. Thus, we propose the CryoDDM-based reconstruction pipeline (Fig. 1), which uses CryoDDM to enhance image quality for more accurate parameters estimation in classification and reconstruction. In this pipeline, we can obtain higher resolution, more conformations, and clearer heterogeneous conformations. Unlike conventional reconstruction pipelines, our approach employs denoised particles not only for particle picking but also for optimizing parameter estimation in subsequent classification and reconstruction, thereby minimizing the impact of noise on the overall reconstruction pipeline. By mitigating noise interference, our pipeline effectively resolves subtle conformational variations and yielding robust results in heterogeneity analysis.

**Fig. 1.**
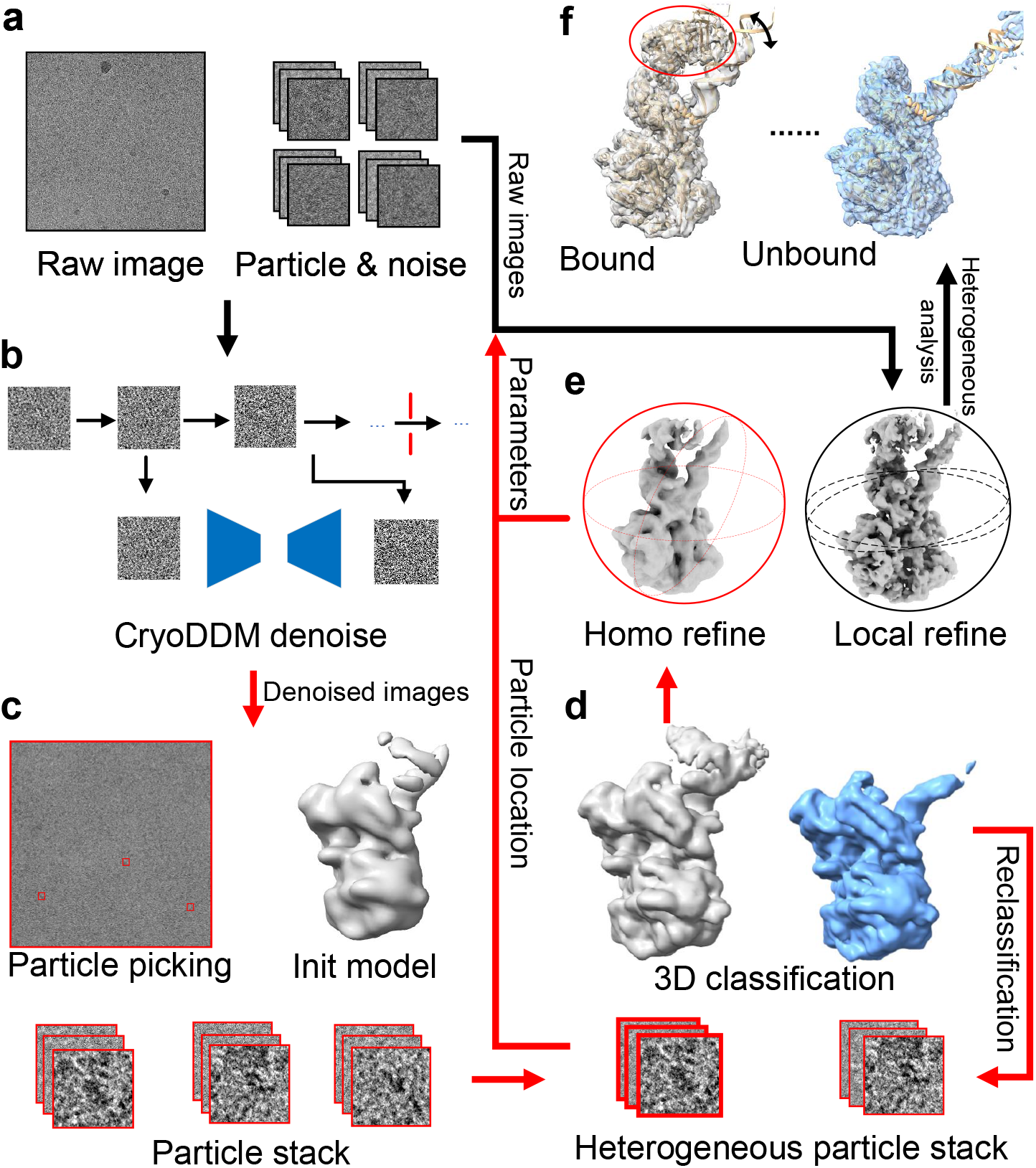
The worflow of CryoDDM-based reconstruction pipeline. **a** Extract particles and noise from raw images to construct the training dataset. **b** Input the data into CryoDDM for denoising training. **c** Denoised images are used for particle picking, 2D classification, and initial reconstruction to enhance the accuracy of these steps. **d** Use denoised particles for 3D classification, and perform reclassification if the initial classification results are suboptimal. **e** Global homogeneous refinement is performed using denoised images for coarse alignment, followed by local refinement with raw images. **f** The classified and corrected 3D volume can be used by any required methods for heterogeneous structural analysis.

As shown in the raw data (black), denoised data (red) in Fig. 1, the proposed pipeline maximizes the use of denoised images to guide protein reconstruction. The raw image is used to extract the particle and noise patch to to the CryoDDM(Fig. 1ab). The denoised images are used to follow traditional pipeline from pick particles to reconstruction initial model(Fig. 1c). If high-resolution results cannot be obtained from the first 3D classification, we can select the dominant classes for multiple rounds of classification to more precisely resolve the structural heterogeneity of the protein(Fig. 1d). Since the 3D classification has already maximally resolved the heterogeneity among protein particles, subsequent structural refinement can be carried out using homogeneous refinement and local refinement(Fig. 1e).

The key to this pipeline is CryoDDM, which preserves protein structural information during denoising, thereby facilitating better particle classification and accurate pose estimation. However, CryoDDM cannot fully distinguish between high-frequency structural information and noise (Fig. 2b), resulting in a slight loss of structural details. Therefore, we use local refinement with raw images to recover these high-frequency details, providing higher resolution 3D structures for subsequent heterogeneous structural analysis tasks.

**Fig. 2.**
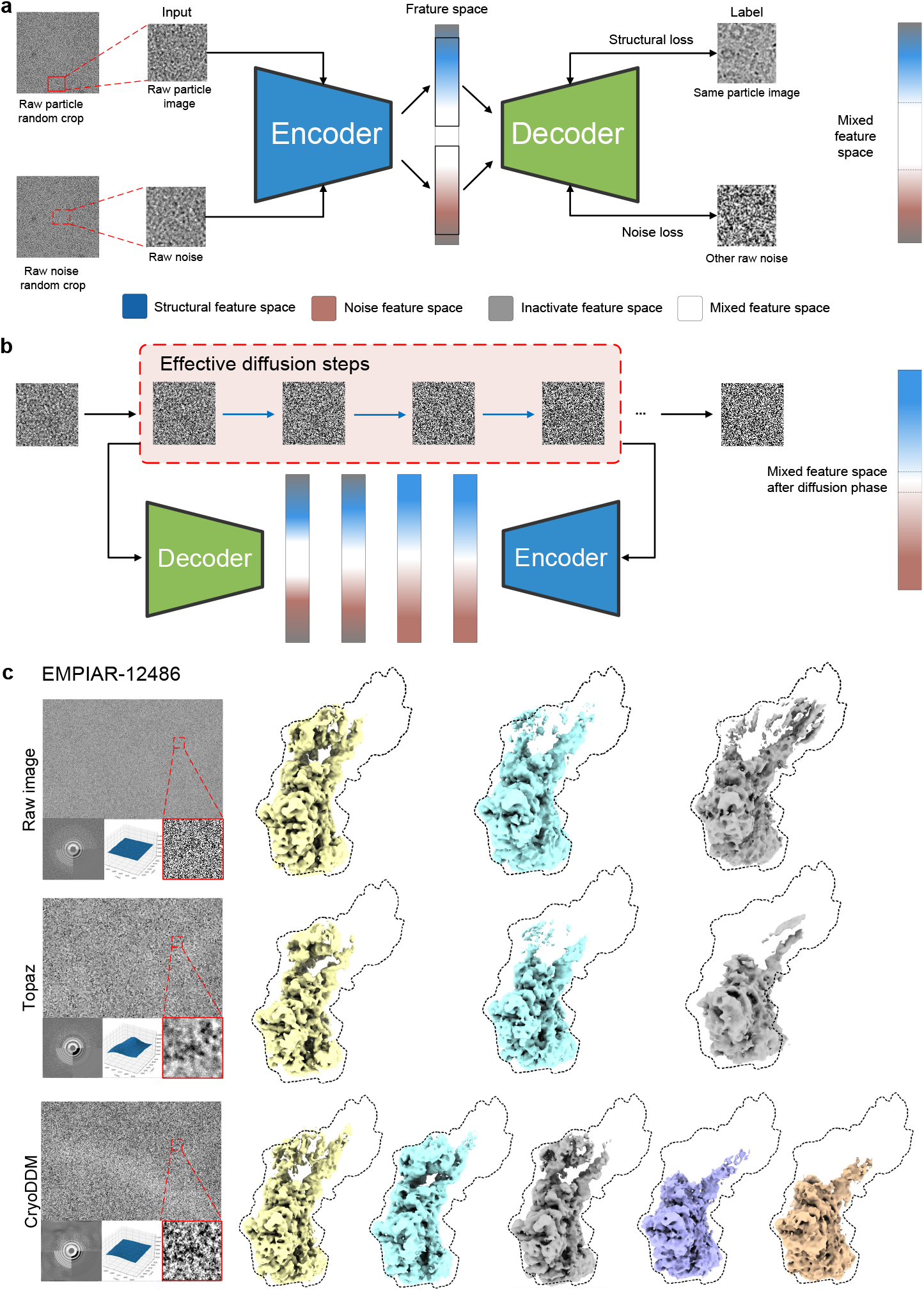
Illustration of the training framework and comparison of denoising methods on 20S proteasome dataset. **a** Crop particle images and noise patches to activate structural feature space and noise feature space. **b** Extract the effective diffusion steps in the diffusion model to achieve further separate noise from structural features while improving the training speed. **c** Denoising performance comparison on EMPIAR-12486. Inset panels reveal that CryoDDM applies non-uniform frequencydomain denoising while maintaining spatial-domain features, whereas Topaz exhibits the inverse behavior (left). This structure feature-preserving property enables CryoDDM to resolve the most conformational states (right).

We validated and compared the results of raw images, Topaz denoised images (for particle picking), the pipeline with Topaz^25^, and the pipeline with Cryo-DDM on multiple datasets (EMPIAR-12087^28^,EMPIAR-10059^29^,EMPIAR-10025^30^, EMPIAR-10345^32^, EMPIAR-12486^38^ and EMPIAR-10827^39^). Our pipeline achieved the best performance (supplementary materials), with significant breakthroughs in heterogeneous conformation classification and reconstruction.

### 2.2 CryoDDM

In recent years, diffusion models have gained significant attention for image denoising tasks due to their exceptional ability to preserve fine details and adaptively reduce noise^40^. Diffusion models often use Gaussian noise to simulate a forward diffusion process, while the noise model of electron microscopy images is typically highly complex and difficult to describe mathematically, making the assumption of Gaussian noise unsuitable for cryo-EM images. Besides, diffusion models typically require sampling over thousands of steps, demanding substantial computational resources, which makes their application in cryo-EM challenging. To address these issues, we propose CryoDDM, a two-phase diffusion model that achieves denoising while preserving structural information. The CryoDDM effectively distinguishes structural information from raw noise through strict guidelines and constraints. Moreover, two-phase CryoDDM enhances the training efficiency, reducing a process that would typically take months or even a year to under 12 hours.

In the first phase, CryoDDM begins by using the same particle image as both the input and the label, which allows it to extract common structural features and activate the structural feature space, as shown in Fig. 2a. Subsequently, CryoDDM uses randomly matched raw noise patches cropped from the same dataset as data pairs for training, enabling the extraction of common noise features and the activation of the noise feature space distinct from the structural feature space. Although CryoDDM at this phase already offers some noise reduction by extracting and recovering particle features, it primarily restores common structural features and does not fully preserve structural details (Supplementary Material). The second phase is the diffusion phase, where we propose a raw noise-based diffusion process to analyze and extract effective diffusion steps, enabling the diffusion model to further distinguish structural information from noise, as shown in Fig. 2b. Fig. 2c shows that the CryoDDM-denoised image is significantly clearer than the raw image, while preserving more high-frequency details than Topaz^25^. The CTF correction closely matches the raw image, with a sharp Thon ring and fewer noise points at high frequencies. Compared with the Topaz reconstruction result, CryoDDM preserves the raw particle structural information more accurately and is able to reconstruct a more detailed 3D structure, as confirmed by the final FSC calculation.

### 2.3 Constraints for CryoDDM in differentiating structural and noise information

The goal of the diffusion phase is to guide the neural network in learning the differences between noise and structural information. In this phase, we propose a residual diffusion model that achieves dual-objective optimization for denoising while preserving structural information. Meanwhile, we introduce the residual structural information low-bound constraint(RSILB) and the training structural loss minimization constraint(TSLM), which determine the most effective steps for preserving structural details within the diffusion model. Finally, we employ an optimization algorithm to compute the most effective steps based on these two constraints, simplifying the sampling and training of the diffusion model and achieving a thousandfold speed improvement.

The forward process of the diffusion model is as follows:

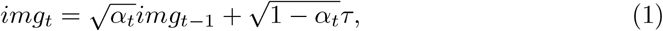

where *t* is the step, *τ* is the noise patch, *α*_*t*_ = 1 *− βt* is a hyperparameter, *img*_*t*_ represents the image at step *t*. In our residual diffusion model, the noise term *img*_*res*_ can be expressed as:

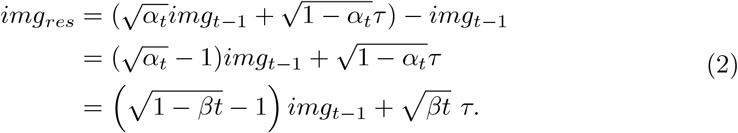

So, we can get the residual diffusion forward process:

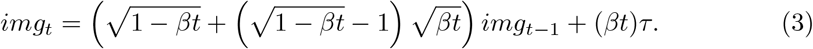

Therefore, the final training objective of our network can be derived as fitting the following function:

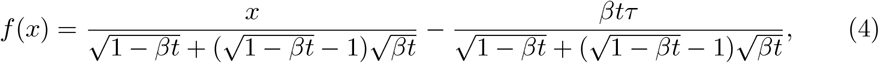

Where 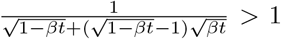 and 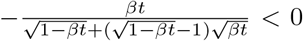, the neural network will tend to enhance the image signal and eliminate noise information. Based on this, we propose the RSILB to reduce the structural features contained in the added noise *τ* :

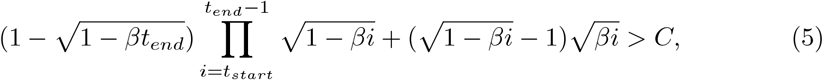

where *t*_*start*_ is the start of forward process, *t*_*end*_ is the end of forward process, *C* is the significant parameter. To ensure that structural information is not lost during the entire training of the diffusion model, we further propose the TSLM:-

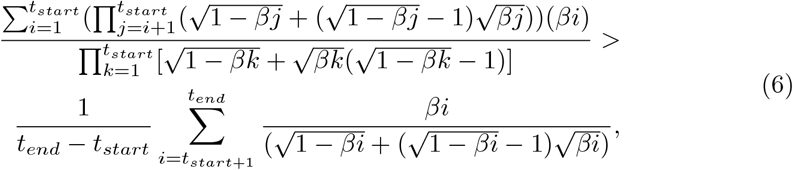

Finally, by maximizing *β* and *t*_*e*_ *−t*_*s*_, we enable fast sampling of the forward process, significantly improving training speed while achieving denoising and preserving structural information.

### 2.4 CryoDDM provides insights into the DNA-protein binding conformation

The core structure of RYBP–PRC1 bound to H2Aub1 mononucleosome adopts a disc-like shape, consisting of a histone octamer wrapped by superhelical DNA. The RYBP–Ub complex associates with one face of the nucleosome through interactions with the acidic patch of histone H2A and the ubiquitin(Ub) moiety, exhibiting a degree of structural flexibility. The RYBP–PRC1 functions as a chromatin-bound signal that promotes the propagation of H2A ubiquitination and reinforces Polycomb-mediated gene silencing. This mononucleosomes with high affinity through cooperative interactions between RYBP’s NZF domain and ubiquitin, and its basic loop with the nucleosome acidic patch, enabling targeted chromatin ubiquitination propagation^41^. The dataset EMPIAR-12087^28^ demonstrates the binding of RYBP to H2Aub1-modified nucleosomes, but does not resolve the dynamic movements of RYBP and Ub. From this dataset, we select 4000 particle images and 150 noise patchs, each with a size of 256 pixels (approximately 1.5 times the particle diameter, 21 nm), where the particle images will be randomly cropped to avoid the phenomenon of centralized attention caused by the training process of the neural network. The training and inference of CryoDDM can be completed within 8 hours on an 4090 GPU.

The denoised images generated by CryoDDM are input into cryoSPARC^18^ for protein structure reconstruction. Following blob and template picking, a total of 1,427,373 two-dimension particle projections are obtained. Subsequently, two rounds of 3D initialization and 3D classification are performed using 5 classes, yielding three distinct conformational states. After homogeneous refinement, local refinement is conducted separately using both the raw and denoised particle stacks. The resulting reconstructions are shown in Fig. 3a. The white structure is reconstructed from the raw particle stack using structural parameters derived from homogeneous refinement of the denoised data, while the blue structure is reconstructed from the denoised particle stack. The overall architecture of the blue structure is correct, but some structural details are lost. The Fourier Shell Correlation (FSC) curves shown in Fig. 3b illustrate the similarity between the two structures, indicating that CryoDDM preserves most of the particle information. Two structures are picked to compare in Fig. 3c. There is a difference in the angle of closure corresponding to the beginning and end of the DNA, while Ub and RYBP also have distinct closure motions. Subsequently, the subunits were separated and fitted into distinct conformational states to investigate the flexibility of the protein. The atomic model of the high-flexibility region is colored red in Fig. 3d. Fig. 3e boxes five regions within the atomic model that exhibit flexible conformational shifts. Region 1 corresponds to Ub, which shows the most pronounced movement, with a tendency to shift toward the histone octamers. Region 2 represents RYBP, whose motion includes a rotational component that appears to correlate with the movement of Ub. Regions 3, 4, and 5 reflect subtle structural variations among proteins within the histone octamers.

**Fig. 3.**
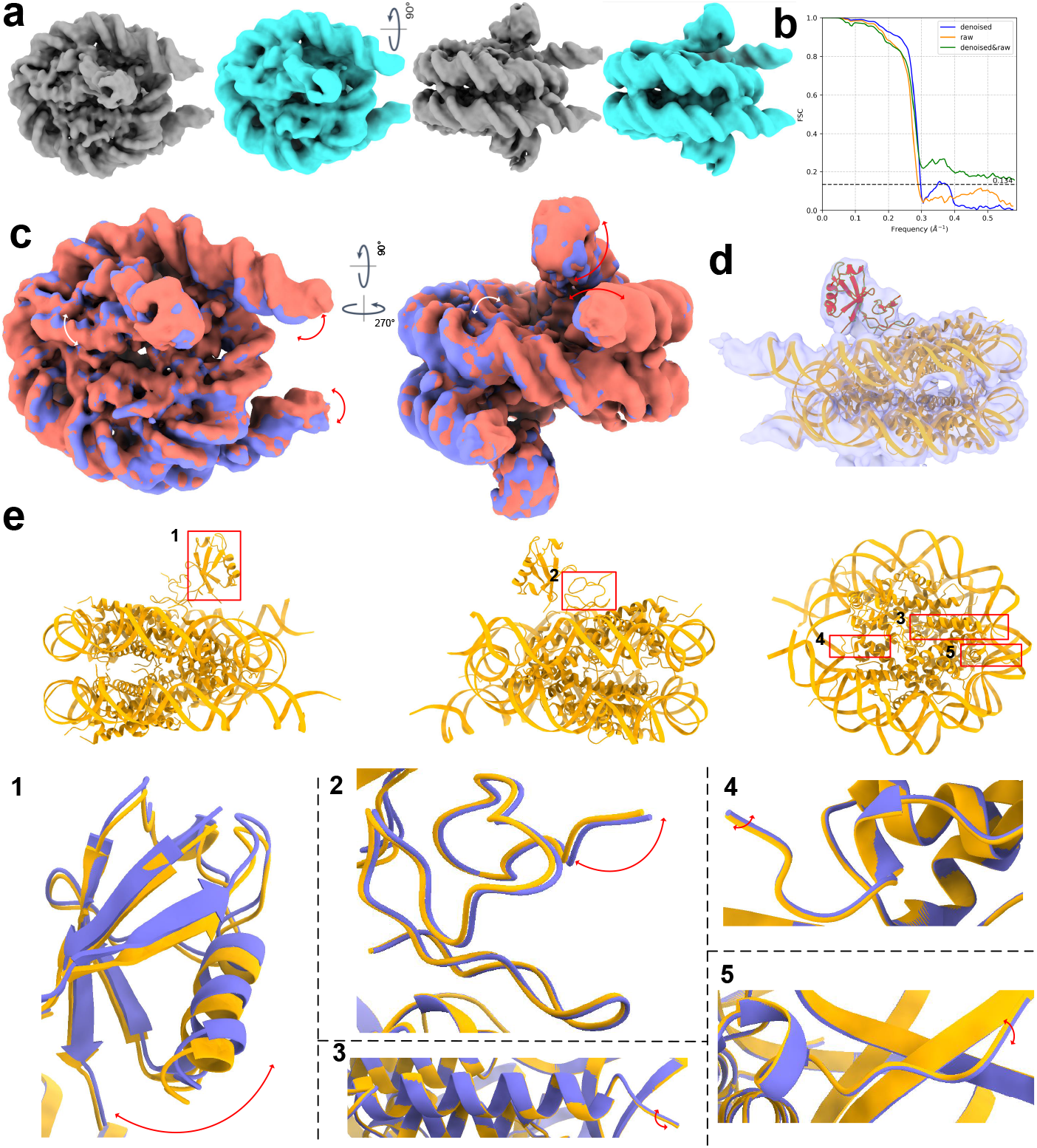
CryoDDM provides insights into the human RYBP-PRC1 bound to H2Aub1 mononucleosome. **a** The reconstruction results from raw and denoised images are compared. The white structure represents the protein model reconstructed from the raw images. The blue structure represents the protein model reconstructed from denoised particles. **b** The FSC curves of 2 structures shown in **a. c** A comparison between conformation 1 (red) and conformation 2 (purple) obtained through the CryoDDM-based reconstruction pipeline. The beginning and end of the DNA shows close movement. The RYBP-Ub region shows up-down movement. **d** Red region represent the most high-flexibility atomic structure. **e** Five flexible region are boxed and enlarged to reveal their movement characteristics. Region 1 is Ub. Region 2 is RYBP. Region 3,4,5 are histone octamers of protein.

### 2.5 CryoDDM uncovers the flexibility of the channel protein

We further explore the application of CryoDDM on a high-symmetry flexible protein. The TRPV1 ion channel, a 380 kDa membrane protein with C4 symmetry, functions as a heat- and capsaicin-activated sensory cation channel. We process the EMPIAR-10059^29^ dataset from raw images, selecting 4000 particles using blob picker and manually choosing 200 background noise patchs for CryoDDM training. The total training completes within 4 hours on an A100 GPU. CTF correction is performed on both raw and CryoDDM-denoised image using CTFFind4^34^, showing no significant changes in the Thon ring, while noise points outside the ring are almost completely eliminated (Supplyment Fig. X).

The CryoDDM denoised images are input into cryoSPARC^18^ for 3D reconstruction. After blob picker and template picker, we obtain 1,183,078 2D projection particle images. Subsequently, we perform 3D initialization and classification with 10 classes, resulting in 6 distinct conformations. Homogeneous refinement is performed using both raw and denoised particles for reconstruction, with the results shown in Fig. 4a. The white structure represents the protein model reconstructed from the raw images, while the blue structure represents the protein model reconstructed from denoised particles. We adjust the visualization levels to the same percentage, revealing the result of the CryoDDM-based reconstruction pipline shows better continuity. Fig. 4b shows resolution of the reconstruction using denoised particles is slightly lower, but the FSC calculations between the two structures demonstrate extremely high similarity in all frequency range (Fig. 4b), confirming that the structure reconstructed from CryoDDM-denoised particles is correct. Next, we export the raw particles for local refinement, and the results are consistent with the structure reconstructed without CryoDDM denoising guidance, as shown in Fig. 4c. The local refinement results show no significant differences: the white structure represents the reconstruction using raw particles, while the blue structure is guided by the denoised homogeneous refinement results. FSC calculations between the structures, presented in Fig. 4d, confirm that CryoDDM-denoised images did not cause reconstruction errors or mislead local refinement. Following the CryoDDM-based reconstruction pipeline, we obtain 6 high-resolution structures, with the main differences shown in Fig. 4e. These include two distinct motions: the first involves all four subunits twisting concentrically around the channel’s pore axis, and the second involves the subunits bends toward or outward, consistent with reports from 3DFlex^12^ and CryoStar^14^. Furthermore, we observe that these two motions often enhance each other, with the amplitude of twisting being positively correlated with the amplitude of rotation. However, unlike those studies, we did not observe significant movement in the basal region of the protein. Besides, we identify the axis of outward rotation for each subunit, where the upper portion exhibits larger movement compared to the lower portion. By fitting the atomic model to different conformations, we identified a red *α*-helix shift (Fig. 4f) that appears to govern the motion of the protein’s upper region. Additionally, we observe some protein motions (Fig. 4g), which is boxed and enlarged to compare with other conformations. For the purpose of maximizing visual contrast, the yellow and purple atomic models may be derived from different conformations. Region 1 shows the *β*-strand shift movement which has been observed by Kwon et al.^42^. Region 2 shows the upper portion bends toward and outward. Region 3 exhibits a red-marked *α*-helix shift, and the flexibility in this region directly contributes to the overall movement of the upper portion of the protein. The *β*-strand shift observed in region 4 appears to be directly driven by conformational changes in region 3.

**Fig. 4.**
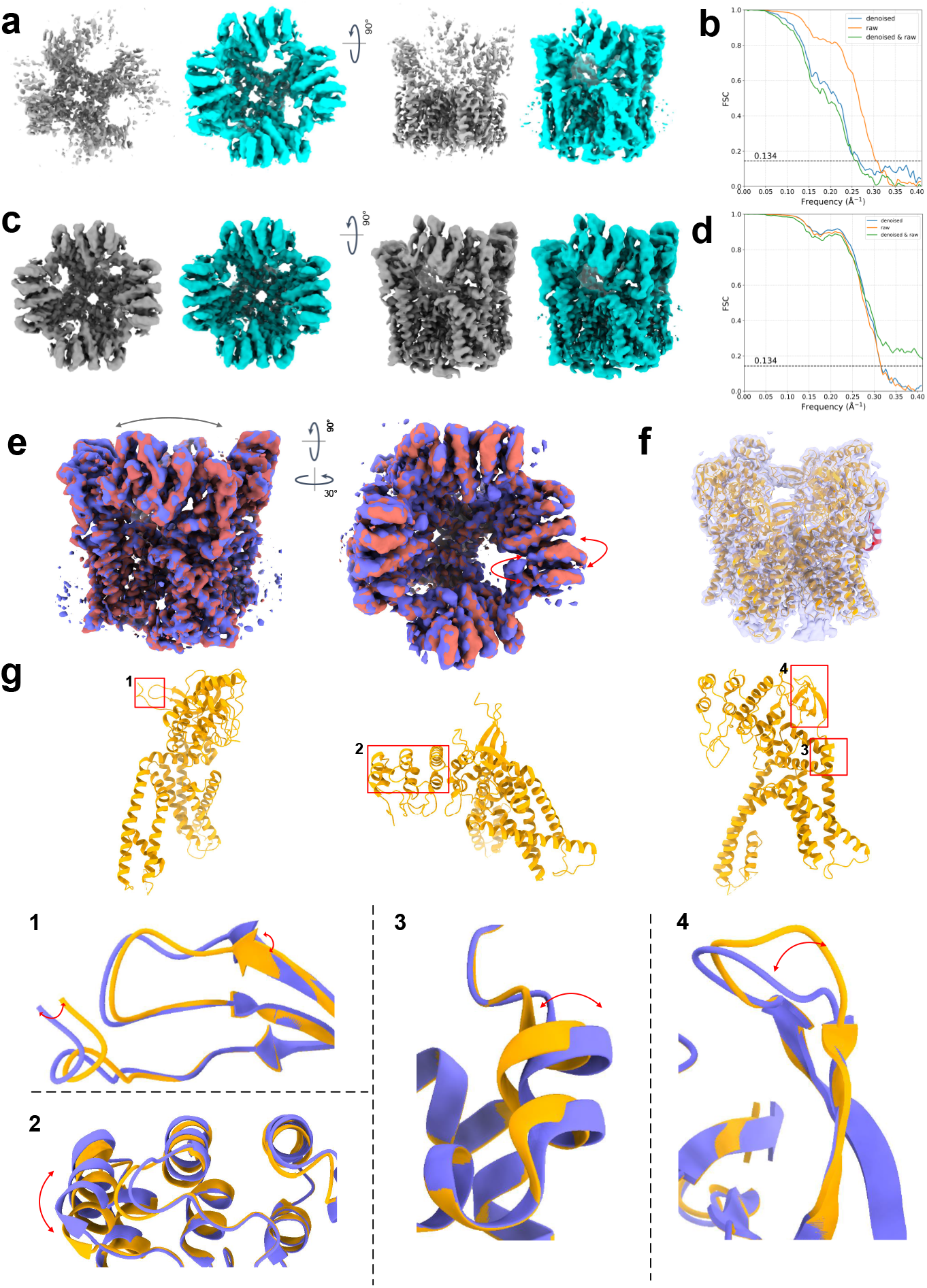
CryoDDM facilitates the classification and reconstruction of multiple dynamic conformations of the TRPV1 ion channel protein. **a** The homogeneous reconstruction results from raw and denoised images are compared. The white structure represents the protein model reconstructed from the raw images and the blue one is reconstructed from denoised particles. **b** The FSC of each protein structure and the FSC between them are shown. **c** Local refinement with or without cryoDDM denoised particle guidance are compared. **d** Reconstruction using denoised-raw particles, raw particles, and their mutual FSC calculations show that the denoised reconstruction is highly similar to the raw reconstruction. **e** A comparison between conformation 1 (red) and conformation 2 (purple) obtained through the CryoDDM-based reconstruction pipeline. **f** The subunit is unstable at the location marked in red, leading to the upper portion structural movement. **g** Take one of the subunit as example, we boxes 4 flexible regions and enlarge them to show the movement.

Compared to other methods, our approach identifies the most valid 2D protein projections (Supplyment Fig. Xa), achieves the highest 3D structural resolution (Supplyment Fig. Xb), uncovers the greatest number of protein conformations (Supplyment Fig. Xc), and exhibits the lowest similarity between these conformations (Supplyment Fig. Xd).

### 2.6 CryoDDM reveals the impact of protein ligands on protein dynamics

To evaluate the performance of CryoDDM in analyzing protein complexes that are prone to local disorder and large-scale conformational movements, we further investigated the structural heterogeneity of the Herpes simplex virus DNA polymerase holoenzyme. It is a double-stranded DNA virus that uses UL30 to replicate the viral genome, while UL42 maintains the stable association of UL30 with DNA^38^. We process the EMPIAR-12486^38^ dataset by selecting 4,000 protein particles using Blob Picker and manually choosing 150 background noise patchs for CryoDDM training. The image pixel size is 256x256, and the total training time was 4 hours. CTF correction is performed on both raw and CryoDDM-denoised image using ctffind4^34^, with minimal impact on the Thon ring, as shown in Supplyment Fig. X.

The CryoDDM denoised images are input into cryoSPARC^18^ for 3D reconstruction. After blob picker and template picker, we obtain 1,192,798 2D projection particle images. Subsequently, we perform 3D initialization and classification with 5 classes, resulting in 2 distinct conformations. By selecting particles from these 2 classes for further 3D classification, 5 structurally distinct protein classes can be obtained. Local refinement is performed using both raw and denoised particles for reconstruction, with the results shown in Fig. 5a. The white structure represents the protein model reconstructed from the raw images,while the blue structure represents the protein model reconstructed from denoised particles. We adjust the visualization levels to the same percentage Fig. 5b shows resolution of the reconstruction using denoised particles is slightly lower, but the FSC calculations between the two structures demonstrate extremely high similarity in all frequency range (Fig. 5b), confirming that the structure reconstructed from CryoDDM-denoised particles is correct. Next, we export the impact of UL42 on protein dynamics. Fig. 5c shows that in the absence of UL42, the DNA exhibits a greater degree of structural mobility, highlighting the crucial role of UL42 in maintaining the stable association between UL30 and DNA. These highly heterogeneous protein atomic structures are highlighted in red in Fig. 5d. Fig. 5e focuses on these two regions, where it can be observed that, in the absence of UL42, the distance between DNA and UL30 increases significantly, and the binding angle becomes markedly larger.

**Fig. 5.**
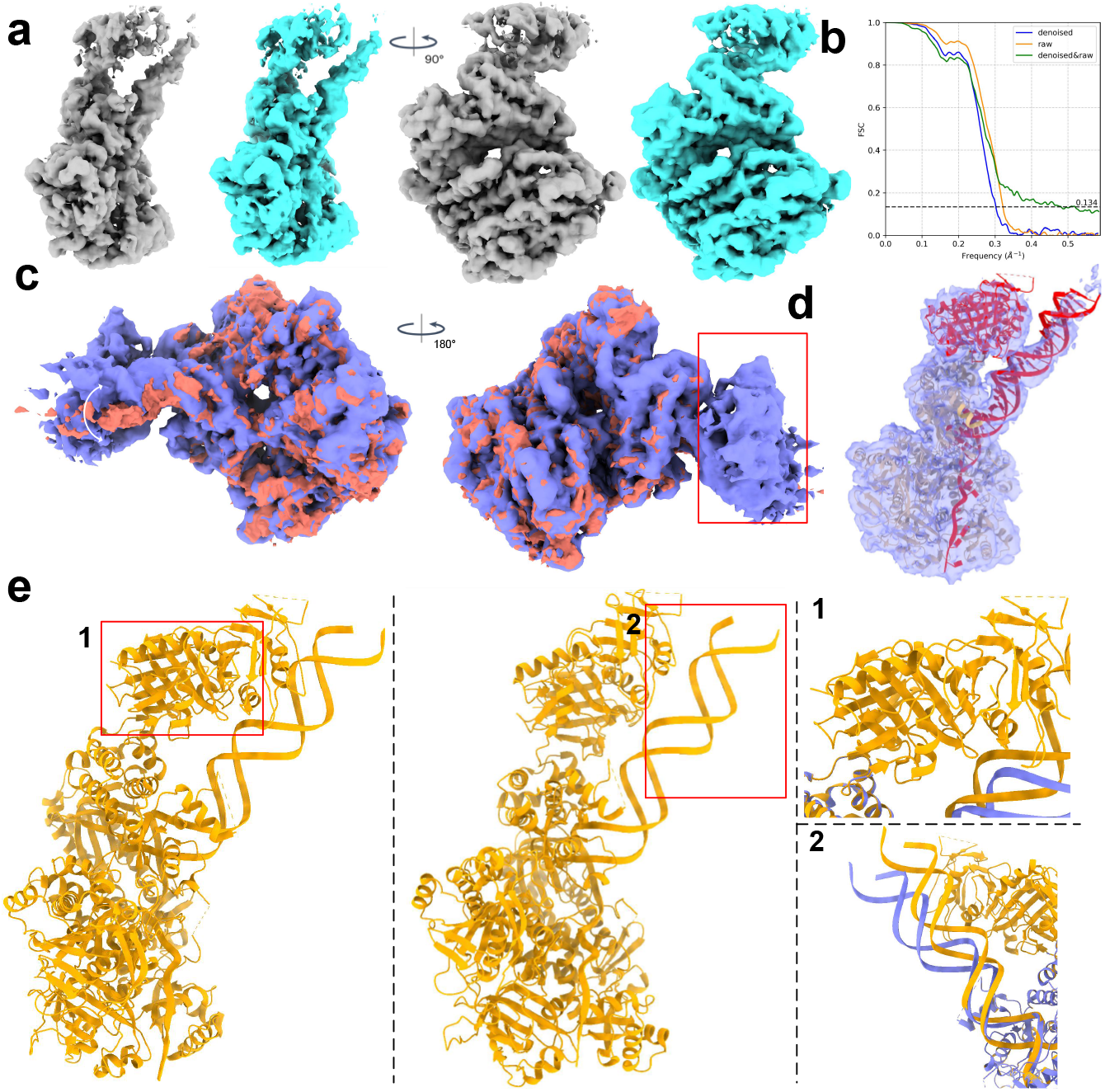
CryoDDM reveals the structural states of the Herpes simplex virus DNA polymerase holoenzyme and DNA under different ligand conditions. **a** The local refinement 3D reconstruction results from raw and denoised images are compared. The white structure represents the protein model reconstructed from the raw images and the blue one is reconstructed from denoised particles. **b** The FSC of each protein structure and the FSC between them are shown. **c** A comparison between conformation 1 (red) and conformation 2 (purple) obtained through the CryoDDM-based reconstruction pipeline. **d** The binding state of UL42 is closely correlated with the state of DNA, and its mobile region is indicated in red in the figure. **e** The two main flexible regions are boxed and enlarged, and the differences in atomic models with and without UL42 binding are shown in yellow and purple.

### 2.7 CryoDDM clarifies dynamic processes of *αV β*8 integrin

To further verify the effectiveness of CryoDDM on highly flexible proteins, we applied it to *αV β*8 integrin. It is a highly flexible protein involved in cell differentiation during development, as well as in fibrotic inflammation and antitumor immunity^32^. We process the EMPIAR-10345 dataset by selecting 4,000 protein particles using Blob Picker and manually choosing 200 background noise patchs for CryoDDM training. The image pixel size is 384x384, and the total training time was 4 hours. CTF correction is performed on both raw and CryoDDM-denoised image using ctffind4^34^, with minimal impact on the Thon ring, as shown in Supplyment Fig. X.

The denoised images are input into CryoSPARC^18^ for reconstruction. After Blob Picker and Template Picker, 73,924 2D projections are selected. The 3D initialization and classification are set to 5 classes, resulting in 5 conformations, three of which have resolutions below 5Å. Raw and denoised partilces are performed with homogenoeus refinement, with the results shown in Fig. 6a, where no significant differences are observed between the two structures. FSC calculations between the structures reconstructed from raw and denoised images (Fig. 6b) demonstrate high similarity, validating the accuracy of the structural information. Both structures are subsequently local refined using raw particle images, leading to further resolution improvement and closer structural similarity (Fig. 6c). This demonstrates that CryoDDM-denoised images can be directly used for protein classification and reconstruction, which is further supported by the FSC curves(Fig. 6d). After that, we obtain 3 high-resolution structures with notable differences, as shown in Fig. 6e. These structures include three major conformational changes: 1) large movements of the protein’s flexible arms, 2) rotation and twisting of the protein’s body structure along the z-axis, and 3) rotation and displacement of the protein’s tail region. Our findings corroborate the work of 3DFlex^12^. Finally, by comparing the CryoDDM-based reconstruction pipeline with the conventional approach, we observe that our method significantly improves reconstruction resolution (Fig. 6f).

**Fig. 6.**
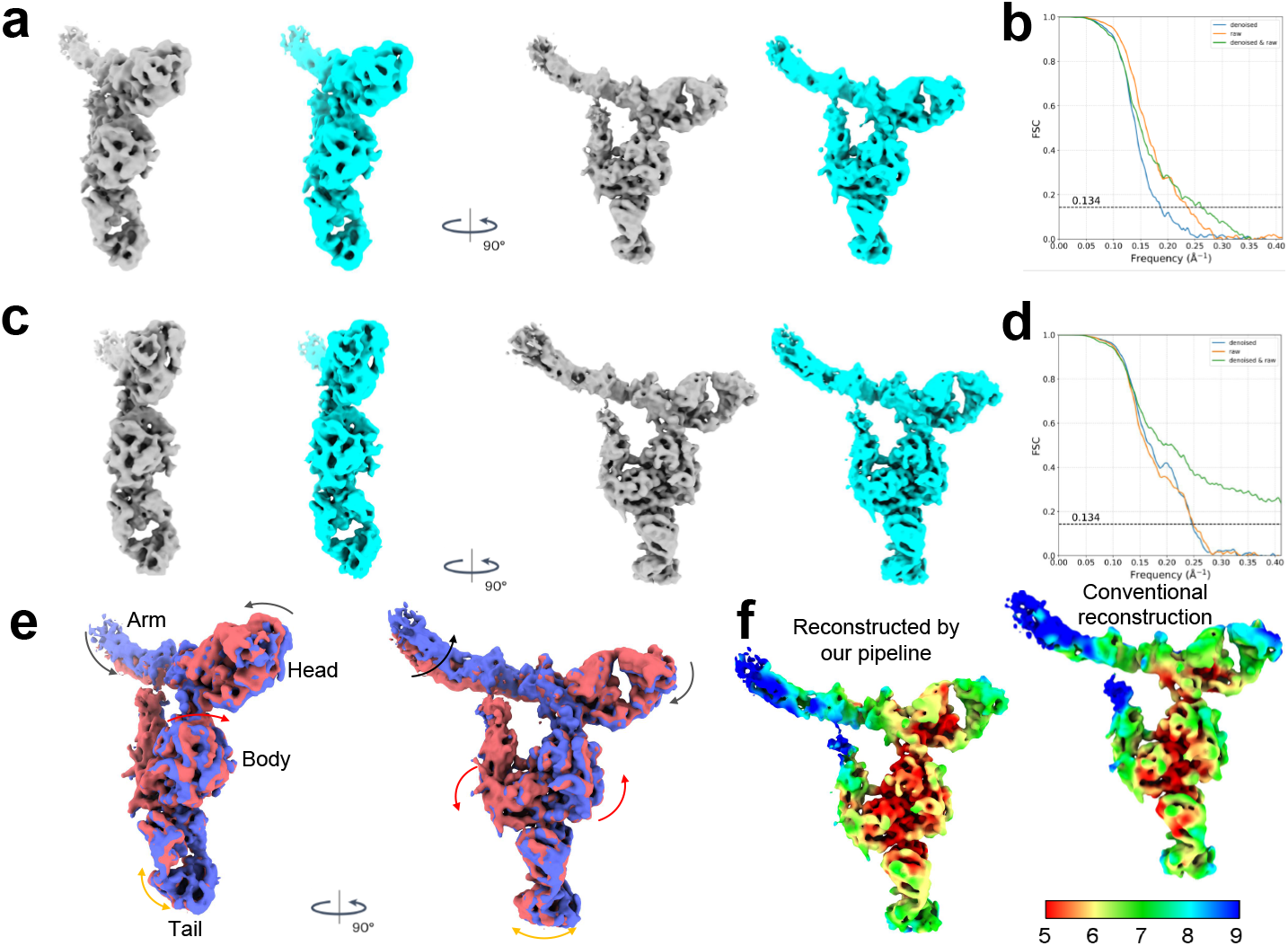
The CryoDDM-based reconstruction pipeline clarifies protein binding and movement processes while improving resolution. **a** Comparison of homogeneous refinement results from the same particle stack between raw and denoised images. The white structure represents the raw image reconstruction, while the blue structure represents the denoised image reconstruction. **b** The FSC curves of each protein structure between them are shown. The resolution of the reconstruction using denoised particles is slightly lower, but it is highly consistent with the raw reconstruction. **c** The local refinement reconstruction is shown, with the white structure representing the raw particles reconstruction and the blue structure corresponding to the denoised results. **d** The FSC curves of each protein structure and between them are shown. After local refinement, the two structures exhibit similar resolutions, and their structural Fourier information is highly similar. **e** The three types of conformational changes of the protein include: the movement of the protein’s flexible arm and head (black arrow), the rotation and twisting of the protein’s body along the central Z-axis (red arrow), and the rotation and twisting of the protein’s tail along the central Y-axis (yellow arrow). Arrows of the same color represent movements belonging to the same group and are interrelated. **f** The CryoDDM-based reconstruction pipeline improves resolution compared to conventional reconstruction pipelines, especially in dynamic regions where the resolution is significantly enhanced.

### 2.8 CryoDDM reveals the motion process of proteasome binding sites

The T20S proteasome is a proteolytic complex responsible for catalyzing protein degradation, with gate opening and closing regulated by the movement of *α* subunits^31, 43^. From the EMPIAR-10025 dataset^30^, we selected 4,000 particle images and 200 noise patchs, each with a size of 384 pixels (approximately 1.5 times the particle diameter, 25.2 *nm*), where the particle images will be randomly cropped to avoid the phenomenon of centralized attention caused by the training process of the neural network. The two-phase training and inference of CryoDDM can be completed within four hours on an A100 GPU.

The CryoDDM denoised images are input into cryoSPARC^18^ for 3D reconstruction. After blob picker and template picker steps, 127,455 2D particle projections are obtained. Subsequently, 3D initialization and 3D classification are performed with five classes, resulting in four distinct conformations. Homogeneous refinement is conducted separately using raw and denoised particles for reconstruction, with the results shown in Fig. 7a. The white structure is reconstructed from the raw particle stack, while the blue structure represents the reconstruction using the same particle stack after Cry-oDDM denoising. No significant structural differences are observed between the two structures in both views. FSC calculations for each and between these structures are presented in Fig. 7b, demonstrating their high similarity in Fourier space. It is shown that our denoising method did not distort the structural information, and the reconstructed particle structures are accurate and reliable. Following this, the raw particle images are exported based on the particle coordinates and subjected to local refinement, yielding four high-resolution structures. The primary difference between these structures lies in the binding site at the top (Fig. 7c), with the movement process shown in Fig. 7d. Detailed structural changes are depicted in Fig. 7e, where the first and fourth panel represent the closed and open conformations, respectively, as identified by Chuah et al^31^. In addition to the two conformations mentioned above, we identified two additional conformations, providing the first explanation of the transition from the closed to the open state.

**Fig. 7:**
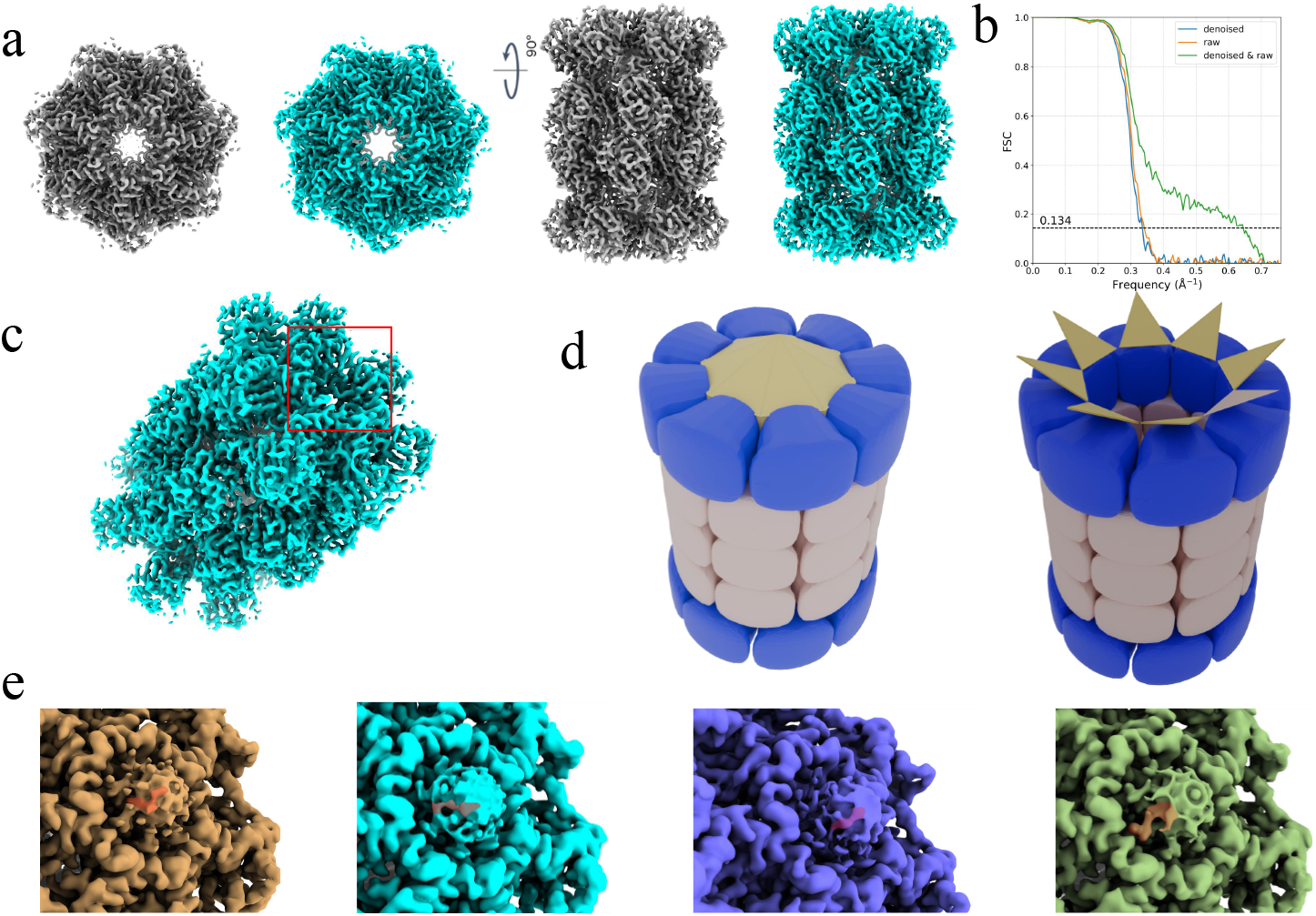
The cryoDDM-based reconstruction pipeline reveals the dynamic process of proteasome binding site movement. **a** Two protein structures from the same particle stack, processed differently, are shown: the white structure is from the raw images, and the blue structure from the CroDDM denoised images. **b** The FSC curves of the two protein structures and between them show are shown. CryoDDM and raw image reconstructions are highly similar in Fourier space, confirming that our denoising model preserves the protein’s structural information. **c** The *α* N-terminus of the 20S proteasome shows a flexible binding site, with its dynamic process displayed in d. **d** Three-dimensional simulations of the T20s proteasome at the closed (left) and open (right) conformational states of its binding site. **e** The structural details of the four protein conformations obtained through classification using the CryoDDM-based reconstruction pipeline are shown. The *α* subunit residues at the binding site, marked by the red mask, gradually rotate during movement and eventually open the gate.

